# Immunogenicity and Protective Efficacy of a SARS-CoV-2 mRNA Vaccine Encoding Secreted Non-Stabilized Spike Protein in Mice

**DOI:** 10.1101/2022.09.07.506878

**Authors:** Eakachai Prompetchara, Chutitorn Ketloy, Mohamad-Gabriel Alameh, Kittipan Tarakhet, Papatsara Kaewpang, Nongnaphat Yostrerat, Patrawadee Pitakpolrat, Supranee Buranapraditkun, Suwimon Manopwisedcharoen, Arunee Thitithanyanont, Anan Jongkaewwattana, Taweewan Hunsawong, Rawiwan Im-Erbsin, Matthew Reed, Wassana Wijagkanalan, Kanitha Patarakul, Tanapat Palaga, Kieu Lam, James Heyes, Drew Weissman, Kiat Ruxrungtham

## Abstract

Establishment of an mRNA vaccine platform in low- and middle-income countries (LMICs) is important to enhance vaccine accessibility and ensure future pandemic preparedness. Here, we describe the preclinical studies of a SARS-CoV-2 mRNA encoding prefusion-unstabilized ectodomain spike protein encapsulated in lipid nanoparticles (LNP) “ChulaCov19”. In BALB/c mice, ChulaCov19 at 0.2, 1, 10, and 30 μg given 2 doses, 21 days apart, elicited robust neutralizing antibody (NAb) and T cells responses in a dose-dependent relationship. The geometric mean titer (GMT) of micro-virus neutralizing (micro-VNT) antibody against wild-type virus was 1,280, 11,762, 54,047, and 62,084, respectively. Higher doses induced better cross-neutralizing antibody against Delta and Omicron variants. This elicited specific immunogenicity was significantly higher than those induced by homologous prime-boost with inactivated (CoronaVac) or viral vector (AZD1222) vaccine. In heterologous prime-boost study, mice primed with either CoronaVac or AZD1222 vaccine and boosted with 5 μg ChulaCov19 generated NAb 7-fold higher against wild-type virus (WT) and was also significantly higher against Omicron (BA.1 and BA.4/5) than homologous CoronaVac or AZD1222 vaccination. AZD1222-prime/mRNA-boost had mean spike-specific IFNγ positive T cells of 3,725 SFC/10^6^ splenocytes, which was significantly higher than all groups except homologous ChulaCov19. Challenge study in human-ACE-2-expressing transgenic mice showed that ChulaCov19 at 1 μg or 10 μg protected mice from COVID-19 symptoms, prevented SARS-CoV-2 viremia, significantly reduced tissue viral load in nasal turbinate, brain, and lung tissues 99.9-100%, and without anamnestic of Ab response which indicated its protective efficacy. ChulaCov19 is therefore a promising mRNA vaccine candidate either as a primary or a boost vaccination and has entered clinical development.

## Introduction

Since COVID-19, the disease caused by severe acute respiratory virus 2 (SARS-CoV-2) began to spread in late December 2019, it has become a pandemic affecting all regions of the world (1). The majority of COVID-19 patients are asymptomatic or only mildly symptomatic (2–4), although the virus is eminently transmissible even during early phases of illness. This contrasts to SARS CoV-1, where peak viral shedding occurred after patients were already quite ill (5, 6). Such unusual characteristics, in conjunction with a highly contagious profile, resulted in rapid spreading of the virus worldwide. In just over 2 years of pandemic, over 10 variants of the virus have been reported, of which 5 (Alpha, Beta, Gamma, Delta, and Omicron) variants have been categorized by WHO as variants of concern (VOCs) (7). These viruses were adapted to increase the transmissibility, severity and/or immune evasion (8). By 18^th^ August 2022, almost 600 million confirmed cases caused by several VOCs and almost 6.5 million deaths were reported (9). Recently, the pandemic is still surging in many countries.

Currently, there are at least 11 approved vaccines using several technology platforms including mRNA, inactivated virus, viral-vectored and recombinant protein (10). The vaccine effectiveness is varied due to several factors such as the emergence of new variants, study population, and prevalence of the outbreak during the period the study was conducted (11–13). Although the currently available vaccines do not completely prevent infection, they are efficacious in reducing severe symptoms in infected individuals (11). Unfortunately, it has also been proven that vaccine efficacy decreases over time (14). Together with the emergence of new VOCs, a booster dose (either homologous or heterologous vaccine modality) is required to enhance the vaccine effectiveness (15).

Among the recently approved vaccines, the mRNA modality seems to be the most efficacious as it induces high levels of desired immune responses and protects from severe symptoms (16, 17). Moreover, the feasibility of large-scale production as well as rapid adaptability to new variants are major advantages of the mRNA production platform. The spike (S) protein of the virus, which contains the major neutralizing epitopes in the receptor binding domain (RBD) and N-terminal domain (NTD), has proved to be the most promising immunogen (18). Thus, most of the recently approved vaccines employ full-length S (with or without modification) or whole virus (inactivated) as a target antigen (19).

Vaccine inequity issue remains a major global challenge. Timely, broad access to effective vaccines in LMICs, particularly the most under-served settings, has always been limited during past pandemics and this has extended to COVID-19 (20). Developing highly effective vaccine platforms like mRNA technology in LMICs is therefore an important goal (21). To respond to COVID19 pandemic and also prepare for future pandemics, Thailand has funded this mRNA vaccine development program from preclinical to manufacturing and clinical development. Here, we describe the construction and preclinical evaluation of mRNA expressing the ectodomain of native, prefusion-non-stabilized S protein of wild-type (WT) strain encapsulated within lipid nanoparticles (ChulaCov19). The vaccine was measured for its immunogenicity in BALB/c mice both using ChulaCov19 alone or in heterologous prime/boost with approved vaccines. It was also evaluated for the protective efficacy in transgenic mice expressing human angiotensin-converting enzyme-2 (ACE2).

## Materials and Methods

### Ethics Statement

The investigators strictly adhered to the principles and guidelines of the Institute of Animals for Scientific Purposes Development, National Research Council of Thailand. All studies were conducted under protocols approved by the Committees on Care of Laboratory Animal Faculty of Medicine, Chulalongkorn University (IACUC approval no. 007/2563), and the Armed Forces Research Institute of Medical Sciences, AFRIMS (IACUC approval no. PN20-06).

### Cells and Viruses

Vero E6, green monkey kidney epithelial cell line was obtained from ATCC (Old Town Manassas, VA, USA). Cells were grown in Eagle’s minimum essential medium (EMEM) supplement with 5% heat-inactivated fetal bovine serum (HIFBS), 1% L-glutamine, 1% Pen/Strep, 40 μg/ml gentamicin and 0.25 μg/ml fungizone (all were from Invitrogen, Carlsbad, CA, USA) at 35±2 °C with 5% CO_2_. One day-old cells were used for measuring of neutralizing antibody by live-virus micro-neutralization (micro-VNT50). The information of SARS-CoV-2 isolates, Wuhan, Alpha, Beta and Delta variants for micro-VNT50 assay performed at the Department of Microbiology, Faculty of Sciences, Mahidol University was described previously (22, 23). For SARS-CoV-2, Wuhan lineage (Hong Kong/VM20001061/2020, NR-52282) that used for micro-VNT50 performed at AFRIMS was obtained through BEI Resources (NIAID, USA). Viruses were propagated in Vero E6 cells to generate sufficient titers 100TCID50 for the micro-VNT50 assay. All isolates were quantitated by tissue culture infectious dose TCID50 using the Reed-Muench method.

### Plasmid Construction

Human codon-optimized sequences of the ectodomain of SARS-CoV-2 spike protein (Wuhan Hu-1 complete genome, GenBank: MN908947.1) was synthesized by GenScript, Piscataway, NJ, USA). It was subcloned into pUC-ccTEV-A101 using *Afe* I and *Spe* I restriction sites (24). The plasmid was propagated in *E. coli* (Stbl3™, Invitrogen, Carlsbad, CA, USA) and extracted by EndoFree^®^ Giga Kit (Qiagen, Hilden, Germany).

### *In vitro* transcription and mRNA encapsulation

Nucleoside modified mRNA was produced by *in vitro* transcription (IVT) by substitution of uridine triphosphate (UTP) with N1-methylpseudouridine (m1Ψ) triphosphate (TriLink, Biotechnologies, San Diego, CA, USA), detailed elsewhere (24). The reaction was carried out employing T7 RNA polymerase (MegaScript, ThermoFisher Scientific, MA, USA) on a linearized plasmid (*Not I/Afl* II double digestions). The mRNA was transcribed to contain 101 nucleotide of adenine (101-poly(A) tails). mRNA capping was performed by the trinucleotide cap1 analog, CleanCap (TriLink Biotechnologies, San Diego, CA, USA). The capped mRNA was purified by cellulose columns purification (25). IVT mRNA was analyzed on agarose for determination of its integrity. Additional quality control to ensure the absence of double stranded RNA (dsRNA) and endotoxin contamination prior to encapsulation into lipid nanoparticles (LNPs) were performed as described previously (26). mRNA encapsulation was performed by Genevant Sciences Corporation (Vancouver, British Columbia, Canada). The proprietary lipid and LNP composition are described in patent application WO2020097540A1 (27, 28). The LNP-encapsulated mRNA were characterized for their size, polydispersity using a Zetasizer (Zetasizer Nano DS, Malvern, UK), encapsulation efficiency, and shipped on dry ice and stored at −80 °C until use.

### mRNA transfection and *in vitro* protein expression analysis

At 24 hr before transfection, 1×10^5^ cells of Vero E6 cells were seeded in 24-well plate (Thermo Fisher Scientific, MA, USA). Cells with approximately 80–90% confluency were transfected with 1 μg of IVT ChulaCov19 using Lipofectamine™ MessengerMax™ (Invitrogen, Carlsbad, CA, USA) according to the manufacture’s protocol. At 24 hr post-transfection, cells were fixed, permeabilized with ice-cold acetone and stained with 1:200 dilution of monoclonal-anti-RBD (R&D Systems, MN, USA), polyclonal-anti-S1-, -anti-S2 antibodies (Sino Biological, Beijing, China), or pooled convalescent serum (PCS) collected in 2020. Goat-anti-mouse IgG-FITC, donkey-anti-rabbit IgG-FITC (both were from BioLegend, CA, USA) or anti-human AlexaFluor647 (Southern Biotech, AL, USA), at dilution of 1:5000 were used as secondary antibodies following anti-RBD, -S1, -S2 or PCS staining. Cell nuclei were counter stained with 4, 6-diamino-2-phenylindole hydrochloride (DAPI) (Sigma-Aldrich, USA). Stained cells were visualized under confocal microscope (ZEISS LSM 800, Carl Zeiss, Germany). For western blot analysis, cell culture supernatant was analyzed by 12% polyacrylamide gel then transferred onto nitrocellulose membrane. The PCS (1:5000) was used for detection of S protein in this step. Goat-anti-human IgG (1:10,000) conjugated with horseradish peroxidase (HRP), (KPL, MD, USA) was used as secondary antibody and detected by chemiluminescence substrate (Immobilon western, Millipore, CA, USA) then exposed to an X-ray film. Recombinant S protein with S1/S2 cleavage site abolished (ACROBioSystems, China) was used as positive control in western blot.

### Immunization in BALB/c mice

To address dose-response study of the ChulaCov19 and heterologous prime-boost responses with other approved COVID-19 vaccines, female BALB/c mice, 4-6 weeks of age, (procured from Nomura Siam International, Bangkok, Thailand) were randomly divided for 5 mice/group into 2 sets of experiment. **Experiment 1**: dose-response of homologous ChulaCov19 prime/boost study, mice were immunized twice intramuscularly with 3 weeks interval of ChulaCov19 with a dosage ranged from 0.2, 1, 10, to 30 μg. **Experiment 2**: a prime/boost regimen of 5 μg of ChulaCov19 and 1/10 of human dosage of approved vaccines available during the study period, including viral-vectored (ChAdOx1; AZD1222, Lot A10062, Nonthaburi, Thailand) and inactivated (CoronaVac, Lot C202105081, Beijing, China) vaccines. In the homologous prime/boost of these 2 approved vaccines groups, each was given in 4 weeks interval. The goal of experiment 2 was to assess the potential role of ChulaCov19 as a booster in a setting of heterologous primed with other COVID-19 vaccine platforms. Mice were bled at 2 weeks after each dose. Antibody responses were measured by ELISA and/or neutralization assays. Splenocytes were collected at 2 weeks after the second dose for assessment of spike-specific IFN-γ T-cell using ELISpot assay (**Figure 1A**).

**Figure 1:**
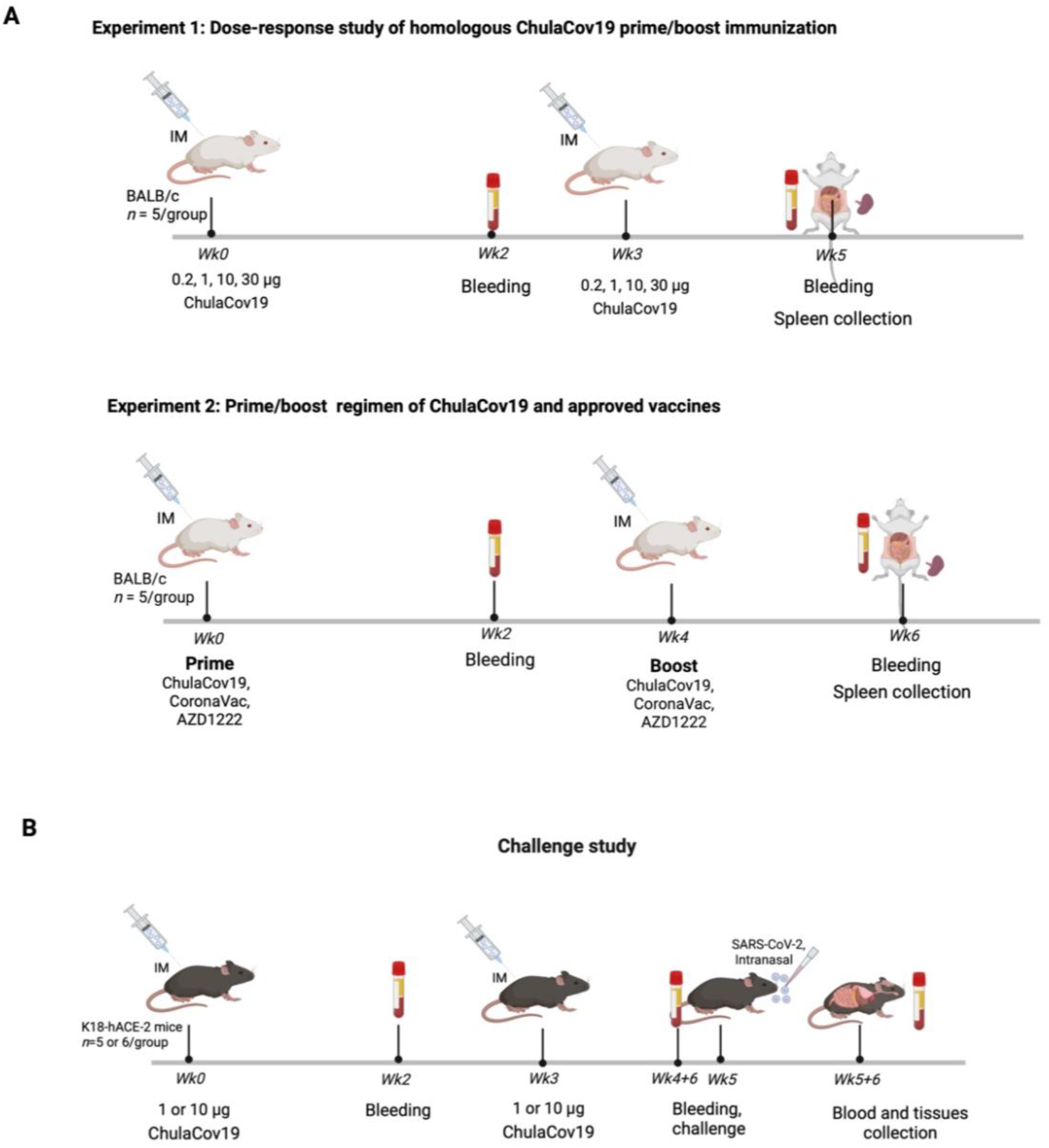
**(A)** Immunization schedule of vaccines in BALB/c mice (*n*=5 per group). ***Experiment 1***: mice were immunized twice intramuscularly (IM) with a 3 weeks interval with various dosages of ChulaCov19 at 0.2, 1, 10 and 30 μg. ***Experiment 2***: For the heterologous prime-boost study, mice were primed with 1/10 of the approved human dosage of CoronaVac or AZD1222 and boosted at 4 weeks later with 5 μg of ChulaCov19. Homologous prime/boost of each vaccine (CoronaVac, AZD1222, or ChulaCov19 were included as control groups. Bleeding was performed at 2 weeks following each dose. Splenocytes were collected at 2 weeks after the second dose. **(B)** Challenge study in K18-hACE2 transgenic mice, *n* = 6 in vaccinated groups and *n* = 5 in control group. Animals were immunized IM with 1 μg or 10 μg of ChulaCov19 at weeks 0 and 3. Sera were collected at weeks 0, 2, 3, 4+6 days, and 5+6 days for NAb measurements. At week 5, mice were challenged intranasally with 2×10^4^ pfu of WT SARS-CoV-2. Tissues were collected at week 5+6 days for assessment of viral RNA. Figures were created with BioRender.com

### Challenge Study in K18-hACE2 Transgenic Mice

Challenge study was conducted under ABSL-3 facility at AFRIMS, Bangkok, Thailand. Seventeen female K18-hACE2 mice (B6.Cg-Tg(K18-hACE2)2Prlmn/J), 7 weeks of age (The Jackson Laboratory, Bar Harbor, ME, USA) were randomly divided into 3 groups. For group 1 and group 2, 6 mice/group were immunized intramuscularly via quadricep muscles with 2 doses, 3 weeks apart of ChulaCov19 at dose of 10 μg and 1 μg, respectively. In negative control (group 3), 5 mice were immunized with PBS instead of ChulaCov19 using the same schedule. At 2 weeks after the second immunization, mice were challenged intranasally with 2×10^4^ pfu (in 50 μL) of SARS-CoV-2 (wild-type). Blood was collected at wk0, wk2, wk3, wk4+6 and wk5+6 days for antibody kinetic analysis **(Figure 1B)**. Six-day post challenge, wk5+6 days, mice were sacrificed for determining virus titers in different tissues (nasal turbinate, brain, lung, and kidney) and for histopathology. Virus titers were quantified by RT-qPCR and by determining the log10TCID50 values.

### Immunogenicity Measurements

#### Total IgG by ELISA

S-specific IgG measurement was performed employing indirect ELISA as described previously (22, 29). In brief, 100 ng of recombinant S-trimer (ACROBioSystems, China) were coated to the 96-well plates. The 5-fold serially diluted mice sera were added in duplicate. After 1 h incubation at 37 °C, plates were washed vigorously with washing buffer (PBS + 0.5% Tween 20). Then, HRP-conjugated secondary antibodies including rabbit anti-mouse IgG (KPL, MD, USA), -IgG1, or -IgG2a (BioLegend, San Diego, CA, USA) were added for an additional 1 h. After washing, the signals were detected by adding tetramethylbenzidine (TMB) substrate (BioLegend, San Diego, CA, USA). The reactions were then stopped with 50 μL of 0.16 N sulfuric acid. The absorbance was measured at a wavelength of 450 nm using a Varioskan microplate reader (ThermoFisher Scientific, Vantaa, Finland). Mid-point titers were calculated and expressed as the reciprocals of the dilution that showed an optical density (OD) at 50% of the maximum value substracted with the background (BSA plus secondary antibody).

#### Neutralizing Antibody

In the immunogenicity dose-response and prime/boost studies (Experiment 1 & 2), NAb measurement was carried out as previously described (22) based on live-virus micro-VNT50 against wild-type, Alpha, Beta and Delta strains in VERO E6 cells with positive cut-off of 1:20. In addition, the pseudovirus neutralization test (psVNT50) against lentiviral pseudovirus bearing a codon-optimized spike gene, described previously (30, 31), was also used for determination of the neutralizing activity against Wild-type, Alpha, Beta, Delta, and Omicron (BA.1 and BA.4/5 subvariants) variants. In the challenge study, NAb was also assessed by live-virus microneutralization test against strain hCoV-19/Hongkong/VM20001061/2020 with slightly different in the incubation period and detection technique, In the latter VNT protocol, serum-virus mixtures were incubated in VERO E6 cells for 5 days. In the detection step, staining of the living cells with 0.02% neutral red (Sigma Aldrich, USA) in 1X PBS (Invitrogen, Carlsbad, CA, USA) was used instead of viral protein staining employing anti-nucleocapsid used in Experiment 1. Lysis solution was added for 1 h at RT before measuring OD at 540 nm. Percentage of virus infectivity in virus control (VC) and samples were calculated based on OD of cell control (CC), infectivity (%) = (OD of CC – OD of sample) x 100. The micro-VNT50 titers was calculated as the reciprocal serum dilution that neutralized 50% of virus observed in virus control wells using probit analysis, SPSS program (32).

#### SARS-CoV-2-spike Specific IFN-γ-producing T-cell Measurement

The procedure of mouse IFN-γ ELISPOT used in this study was described in our previous reports (22, 33). In brief, mouse splenocytes at 5×10^5^ cells/well were cultured with SARS-CoV-2 spike peptide pools spanning the entire sequence of spike protein, 25 peptides/pool (Mimotopes, Mulgrave, Victoria, Australia) at a final concentration of 2 μg/mL at 37 °C, 5% CO_2_ for 40 h. Pools 1–5 and 6–10 corresponded to S1 and S2 regions of spike protein, respectively. Secreted mouse IFN-γ was captured by anti-mouse IFN-γ (AN18) monoclonal antibody (Mabtech, Nacka Strand, Sweden) precoated on 96-well nitrocellulose membrane plates (Merk Millipore, Darmstadt, Germany). Results were expressed as spot-forming cells (SFCs)/10^6^ splenocytes after subtraction of the spots from negative control wells.

### Real-time PCR for Quantification of SARS-CoV-2 RNA

SARS-CoV-2 RNA levels in serum and tissue samples were quantitated using quantitative RT-PCR. Viral RNA was extracted from 140 μl serum and tissue samples using the QIAamp viral RNA mini kit (QIAGEN, Hilden, Germany). For tissue samples, RNeasy Mini Kit (QIAGEN, Hilden, Germany) was used following the manufacturer’s instruction. The total volume of 50 μl of viral RNA was obtained from each sample. Five microliters of each RNA sample was used in quantitative RT-PCR that was performed using CDC procedure (34) and AFRIMS SOPs *in vitro* SARS-CoV-2 RNA transcripts (IVTs). In each experiment, 3 internal controls (No Template Control (NTC), Negative Extraction Control (NEC) and Positive Extraction Control (PEC)) and 6 *in vitro* transcribed RNA standards were run along with test samples in each experiment. The number of copies of viral RNA per sample was derived from standard curves of serial dilutions of IVTs (5, 50, 5×10^2^, 5×10^3^, 5×10^4^, 5×10^5^ RNA copies number or genomic equivalent (GE)/reaction were included. The GE/ml of virus in a serum sample was calculated by multiplying the number of copies/reaction by [10,000 x the volume of a serum sample used (μl) for extraction]. The GE per gram of virus in a tissue sample was calculated by multiplying the number of copies/reaction by [10,000 x the weight of a tissue sample (mg) used for extraction].

### Histological study

After euthanasia, tissue samples were excised and fixed in 10% formalin. After fixation, samples were demineralized for 24 h using Decal Stat (Decal Chemical Corp., NY, USA), rinsed with sterile deionized water for 3-5 min, trimmed in a dorsal–ventral plane bisecting the spinal column, and placed back into 10% formalin. Tissue specimens were embedded in paraffin, thin-sectioned, mounted on positively charged glass slides, and stained with hematoxylin & eosin or anti-SARS-CoV-2 antibody for immunofluorescent assay.

### Statistical analysis

Statistical analysis was performed using GraphPad Prism 9.0 software (San Diego, CA, USA). Comparisons of the data between groups were made using non-parametric t tests (Mann– Whitney or Wilcoxon signed rank tests). All *p* values < 0.05 were defined as statistically significant.

## Results

### *In vitro* Protein Expression

The purified mRNA-S (ChulaCov19) with undetectable endotoxin was tested for protein expression in VERO E6 cells. By using immunofluorescent assay, employing RBD-, S1-, S2-specific antibodies or PCS, S proteins were observed within the cytoplasm of transfected cells while transfected cells were negative for fluorescent signal **(Figure 2A)**. By using western blot, S protein could be detected in cell culture supernatant when using PCS as primary antibody. The bands corresponded to S1, S2 and also intact S. Comparable molecular weight of the intact S expressed by ChulaCov19 was also observed when used commercial recombinant S with S1/S2 cleavage site abolished as a control.

**Figure 2:**
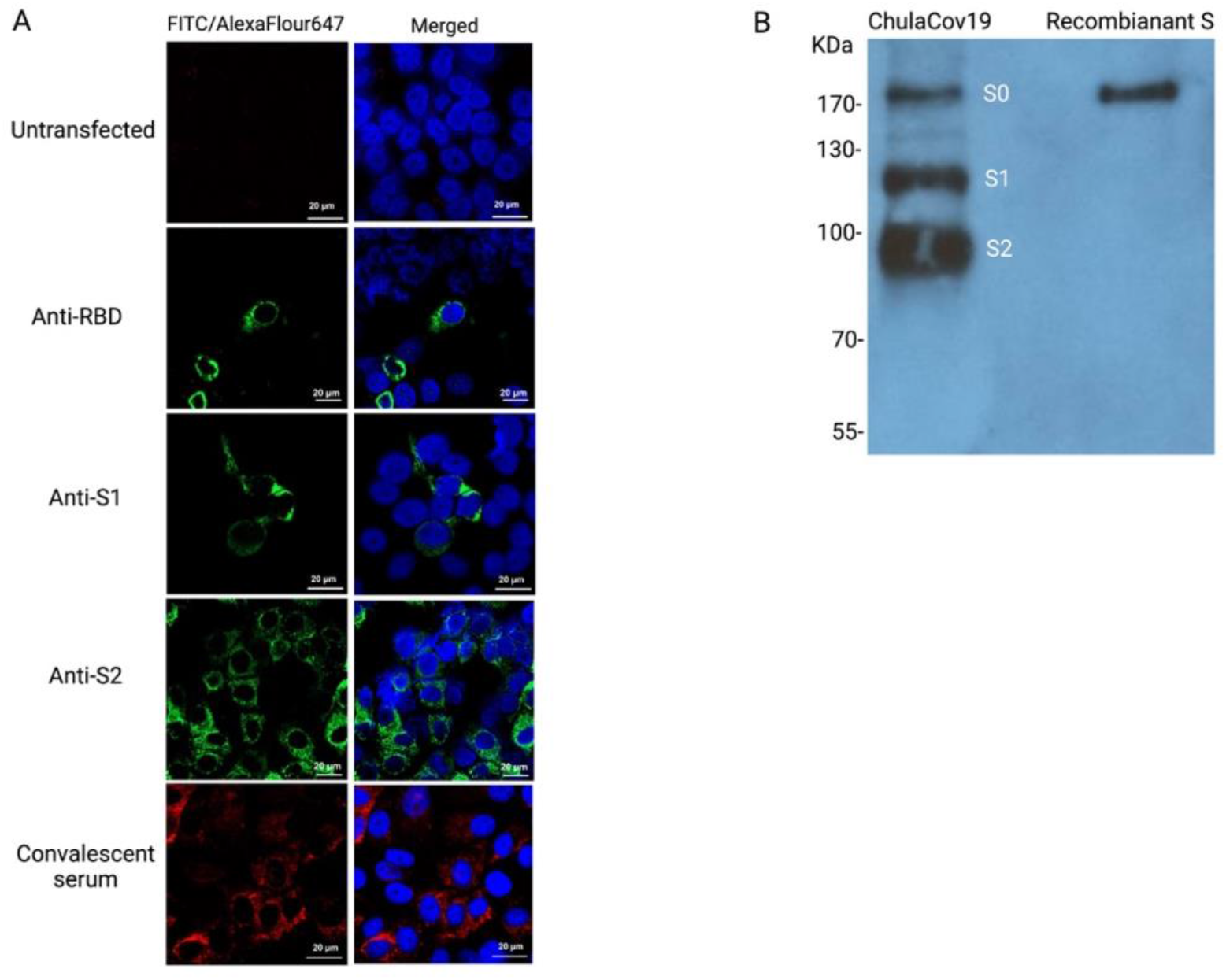
Protein expression analysis after 24 h transfection of ChulaCov19 in VERO E6 cells. **(A)** Intracellular S protein expression examined by immunofluorescent assay employing anti-RBD, -S1, -S2 or PCS as primary antibody, the nuclei were counter stained with DAPI (blue). FITC-tagged 2^nd^ Abs (green) were used for detection of RBD, S1, and S2 while AlexaFluor647-tagged 2^nd^ Ab (red) was used following PCS staining. **(B)** S protein expression in cell culture supernatant analyzed by western blot. Recombinant S protein (10 ng) with abolished S1/S2 cleavage site loaded in the right lane was used as positive control.

### Spike-specific Total IgG Antibody Analysis of One versus Two Doses of Immunization

The S-specific total IgG after 1 or 2 doses of ChulaCov19 was analyzed in mice sera from experiment 1. S-specific total IgG analyzed at week 2 revealed that all ChulaCov19-immunized mice, either with 1 or 2 doses, elicited anti-S-specific IgG response started at the lowest dose of 0.2 μg with a dose-dependent response pattern. The second dose of ChulaCov19 strongly augmented the IgG antibody levels with an increase of 10-19 folds **(Figure 3A)**. IgG1 and IgG2a subclasses were also assessed to determine Th2 and Th1 responses, respectively. The results demonstrated that IgG2a/IgG1 (or Th1/Th2) ratios were greater than 1 in all vaccinated mice **(Figure 3B)**. These results reflecting that ChulaCov19 was highly immunogenic and induced a Th1-skewed response in mice.

**Figure 3:**
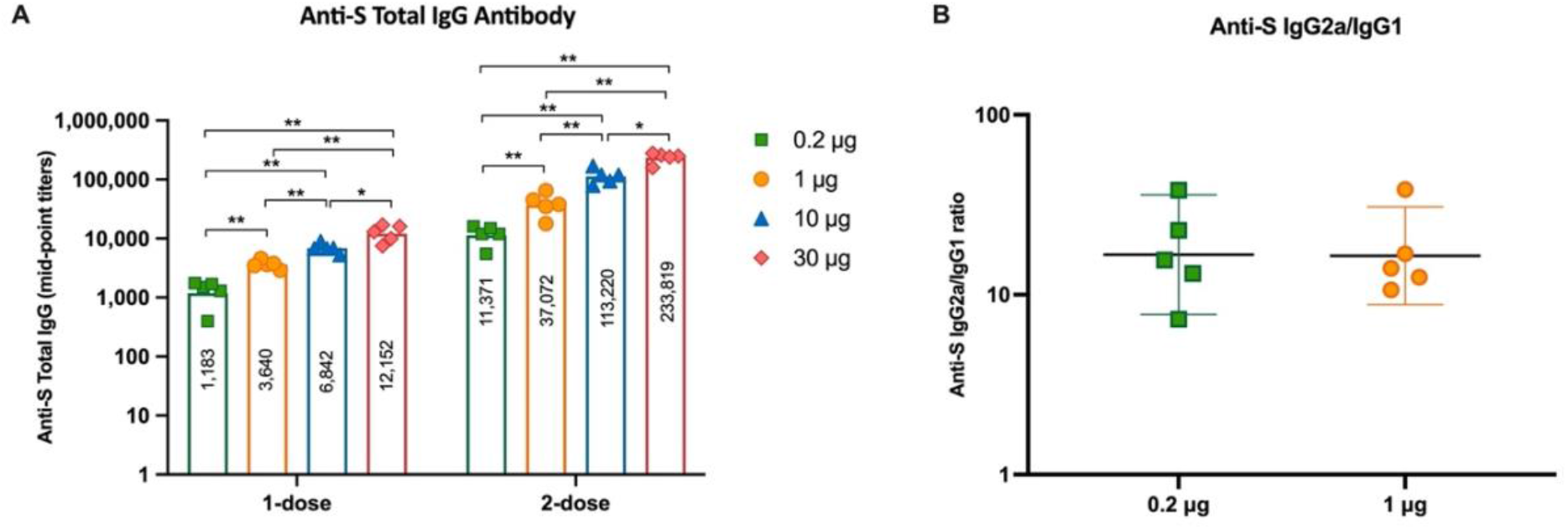
S-specific IgG responses in ChulaCov19 immunized mice. (A) Kinetics of total IgG at 2 weeks after receiving 1 or 2 doses of 0.2, 1, 10, and 30 μg of ChulaCov19. (B) S-specific IgG2a/IgG1 ratio measured at 2 weeks after the 2^nd^ dose. (Note: the IgG2a/IgG1 ratio of 10 μg and 30 μg immunized mice were not analyzed due to limited volume of serum samples). Bars (A) or horizontal lines (B) represent the geometric mean (GMT) for each group while error bars indicate the 95%confident interval. Statistical analysis significance was determined by Mann–Whitney test. P<0.05 and P<0.01 are indicated by * and **, respectively.

### Neutralizing Antibody Results

NAb measurement in mice sera from Experiment 1 against WT live-virus (micro-VNT50) at 2-week after received each dose showed NAb response in a dose-dependent manner. After the first dose, NAb were detected in mice that were received 1, 10, and 30 μg ChulaCov19 with GMTs of micro-VNT50 titer of 80, 368, and 735, respectively. The NAb titers were drastically enhanced after the second dose was given. After 2-dose, the GMTs of MN50 titer for 0.2, 1, 10, and 30 μg were 1,280, 11,763, 54,047, and 62,084, respectively **(Figure 4A)**. This finding implied that ChulaCov19 is highly immunogenic against wild-type strain. Mice sera were further analyzed for NAb by psVNT50 test against the important recent VOCs, including Delta and Omicron (BA.1, and BA.4/5) variants, and titers were significantly decreased for all VOCs. For example, at 10 μg dosed group, the GMTs of psVNT50 for Delta and Omicron (BA.1) variants were decreased for 5.9 and 14.3 folds when compared against wild-type strain **(Figure 4B)**. psVNT50 against BA.4/5 subvariant showed the lowest GMT in all dose ranges. When compared with psVNT50 titers against BA.1, the GMT fold reduction against BA.4/5 in 10 and 30 μg dosed groups were 48 and 2.3 folds, respectively. This result implied that the decrease in NAb titers against BA.4/5 may be improved with higher the mRNA vaccine dose.

**Figure 4:**
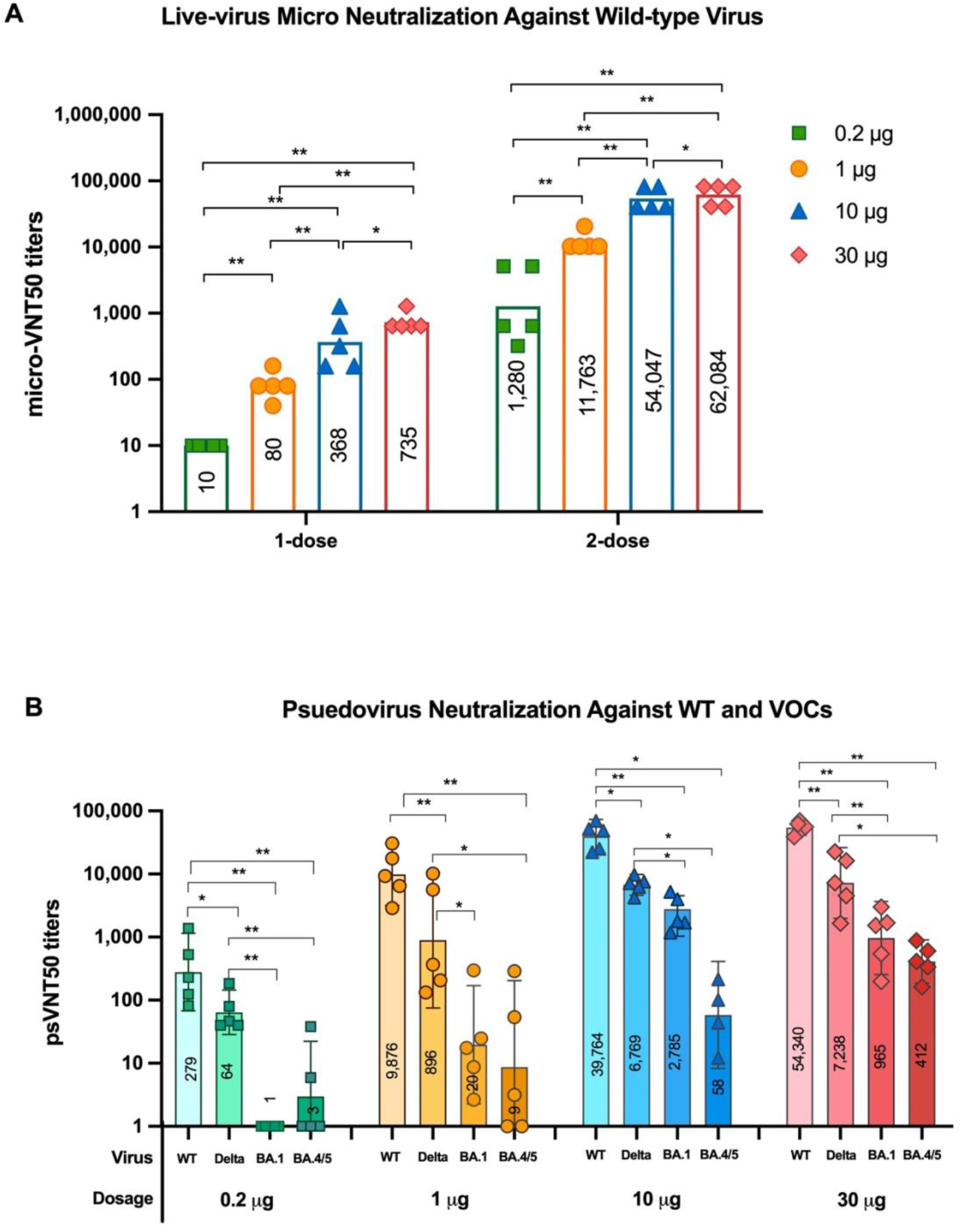
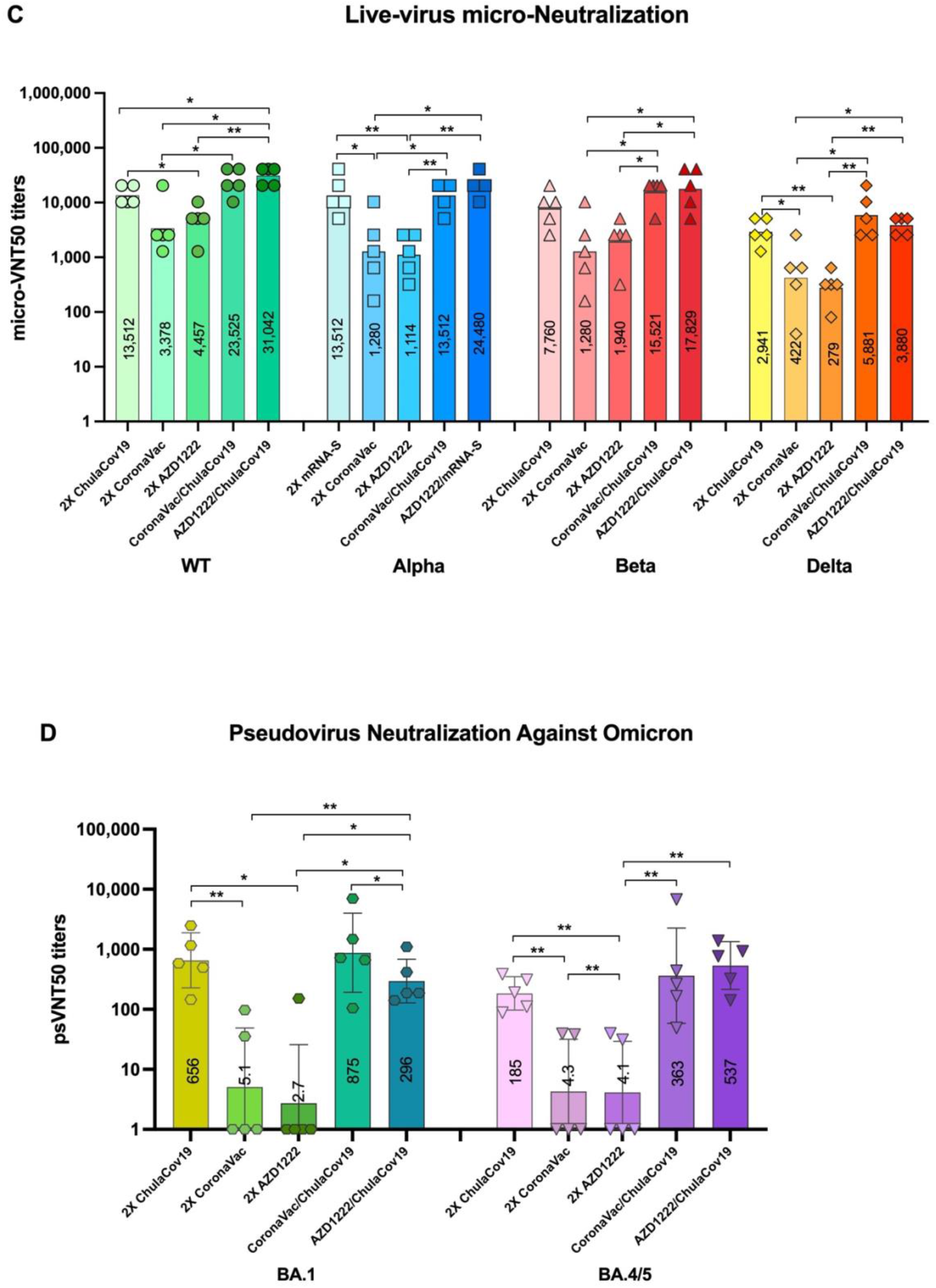
NAb responses in immunized mice. ***Experiment* 1: (A)** Live-virus microneutralization (micro-VNT50) titers against WT live-virus at two weeks after receiving each vaccine dose. **(B)** Pseudovirus neutralization test (psVNT50) titers at two weeks after the second dose againt WT, Delta, Omciron (BA.1, and BA.4/5) variants. ***Experiment* 2**: **(C)** micro-VNT50 titers against WT, Alpha, Beta, and Delta live-virus at two weeks after receiving various homologous or heterologous prime/boost regimens. **(D)** psVNT50 NAb titer results at two weeks after the second dose in various prime/boost regimens againt Omicron BA.1 and BA.4/5 subvariants. Each bar represents the GMTs and 95% CI for each group. Statistical analysis significance was determined by Mann–Whitney test. P<0.05 and P<0.01 are indicated by * and **, respectively.

ChulaCov19 was further compared to two approved vaccines CoronaVac or AZD1222, either in a homologous prime/boost setting or heterologous one (i.e. as a booster dose in mice that had been primed with CoronaVac or AZD1222 **(Experiment 2)**. At the same dose range which is 1/10 of human dosage of approved vaccines and 5 μg of ChulaCov19, in the homologous prime/boost comparison, ChulaCov19 showed 3-to 10.6-fold higher NAb level than 2-dose immunization of CoronaVac or AZD1222 across all variants (Wild-type, Alpha, Beta, and Delta), as measured by micro-VNT50 **(Figure 4C)**. In the heterologous prime/boost comparison, mice primed with CoronaVac or AZD1222 and boosted with ChulaCov19 generated significantly higher GMT against WT, Alpha, Beta, Delta, and Omicron than the respective homologous prime/boost groups but were comparable to that of the ChulaCov19 homologous prime/boost. For example, the micro-VNT50 GMT against WT in the AZD1222-prime/ChulaCov19-boost group was 7-fold higher than 2-dose AZD1222 immunization (GMT of micro-VNT50 were 31,042 *vs* 4,457, *p* = 0.0079). And the GMT NAb titer against WT in the CoronaVac-prime/ChulaCov19-boost group was also 7-fold higher than 2-dose of the CoronaVac group (GMT of micro-VNT50 were 23,525 *vs* 3,378, *p* = 0.0317), **Figure 4C**. In the case of Omicron variants, psVNT50 NAb GMT results against Omicron BA.1 and BA.4/5 subvariants showed similar findings **(Figure 4D)**. Although the neutralization against those VOCs were significantly reduced when compared with WT, the heterologous prime/boost regimen was more efficient (84-172 folds increase) in inducing cross-NAb against VOCs including BA.1 and BA.4/5 subvariants than homologous CoronaVac or AZD1222 immunization **(Figure 4C, 4D)**. These results confirmed that ChulaCov19 is highly immunogenic either as a primary vaccination in a vaccine naïve setting, or as a booster vaccine in animals previously vaccinated with the other vaccines.

### SARS-Cov-2-spike Specific T cell Responses

Splenocytes from mice immunized with various dosages of ChulaCov19 (Experiment 1) were analyzed as summed frequency of S-specific IFN-γ positive T cells (Figure 5A). Similar to the antibody results, the magnitude of T cell response was found to be dose-dependent but peaking at the 10-μg dosage. However, the slightly higher level compared to the 30-μg group was not statistically significant. Mean spike-specific IFN-γ positive T cells for 0.2, 1, 10 and 30 μg were 166, 429, 1,913, and 1,378 SFC/10^6^ splenocytes, respectively. T-cell responded to S1-pooled peptides much more common than to S2-pooled peptides. The analysis of the responses to different part of S-specific pool peptides in all vaccinated groups showed that peptide pool #3-5 (which include receptor-binding domain or RBD) and pool #9 (which includes Heptad Repeat 2 or HR2) in S1 and S2, respectively, were the most common peptides pools that recognized by the vaccinated mice T-cells **(Figure 5A)**.

**Figure 5:**
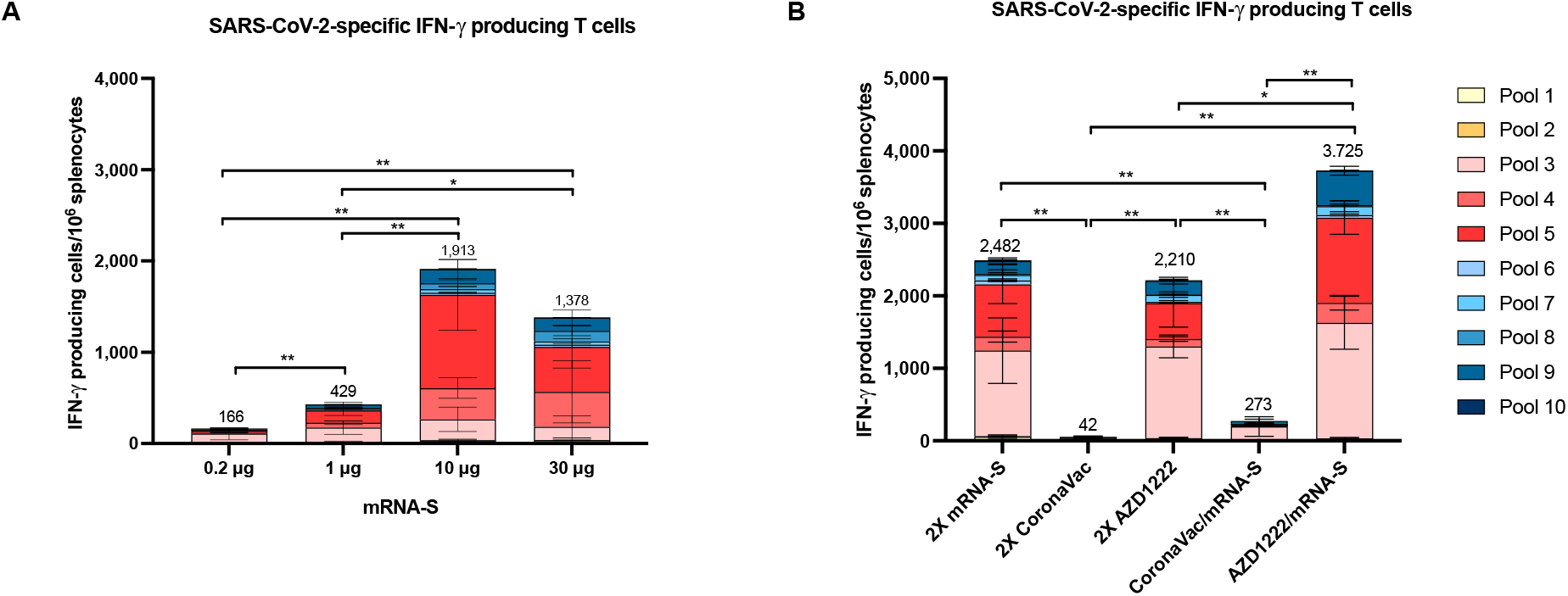
Induction of S-specific IFN-γ positive T cells in BALB/c mice immunized with various doses of ChulaCov19 analyzed at 2 weeks after the second dose **(A),**and in heterologous prime/boost study in mice primed with with CoronaVac or AZD1222 vaccine and boosted with ChulaCov19 (5 μg). Homologouse prime/boost results of each vaccine were included. Bars represent the mean ± SD of S-specific IFN-γ positive T cells after stimulated with overlapping peptide pools spanning the SARS-CoV-2 S1 (pooled #1-5) and S2 (pooled #6-10). Statistical analysis significance was determined by Mann–Whitney test. P<0.05 and P<0.01 are indicated by * and **, respectively.

In the heterologous *vs* homologous prime/boost experiment (Experiment 2), homologous ChulaCov19 and homologous AZD1222 immunizations elicited comparable levels of S-specific IFN-γ positive T cells responses which was 2,482 and 1,899 SFC/10^6^ splenocytes, respectively. Of interest, the heterologous AZD1222-prime/ChulaCov19-boost induced the best specific T cells responses with mean spike-specific IFN-γ positive T cells of 3,725 SFC/10^6^ splenocytes, which significantly higher than all groups except homologous ChulaCov19. In contrast, CoronaVac immunization showed the lowest T cells responses (42 SFC/10^6^ splenocytes). Boosting with ChulaCov19 significantly enhanced the magnitude of T cells response in CoronaVac-primed mice (273 SFC/10^6^ splenocytes) but it still far lower than using homologous ChulaCov19 or AZD1222-prime/ChulaCov19-boost immunization regimens **(Figure 5B)**.

### Protective Efficacy of ChulaCov19 in K18-hACE2 Transgenic Mice

#### Immunogenicity results

After received 2 doses of ChulaCov19 or phosphate-buffered saline (PBS, control group) 3 weeks apart, K18-hACE2 mice were tested for NAb kinetics against live SARS-CoV-2 strain hCoV-19/Hongkong/VM20001061/2020. Baseline NAb levels at week 0 of all mice were negative. However, at week 2 after the first dose, 6/6 and 4/6 animals from the 10 μg and 1 μg groups, respectively, showed a dose dependent manner of NAb response to vaccine administration with significant difference in GMTs of micro-VNT50 titers (*p* = 0.0065). At week 3 after dose 1, NAb were still detected in all animals in the 10 μg group, and 5/6 animals in the 1 μg group. At week 5 (2 weeks after the second dose), all mice in both vaccinated groups showed a significant increase of NAb levels with GMT of micro-VNT50 titers of 15,343 and 4,424 in the 10 μg and 1 μg group, respectively, *p* = 0.0325. Day 6 after the viral challenge (Week 5 +6 days), there was a slight decline of NAb titers in both groups. In contrast, sham-treated animals failed to show any NAb response except for one animal on Wk5+6d (**Figure 6A**). Of note, all vaccinated mice at the 10 μg dose, and 5 of 6 mice at 1 μg dose elicited SARS-CoV-2 specific serum IgA (data not shown). There was no anamnestic response (four-fold increase on MN50 titers) in all vaccinated group 6 days after the challenge, whereas one mouse in the control group developed low micro-VNT50 titer at 40.

**Figure 6:**
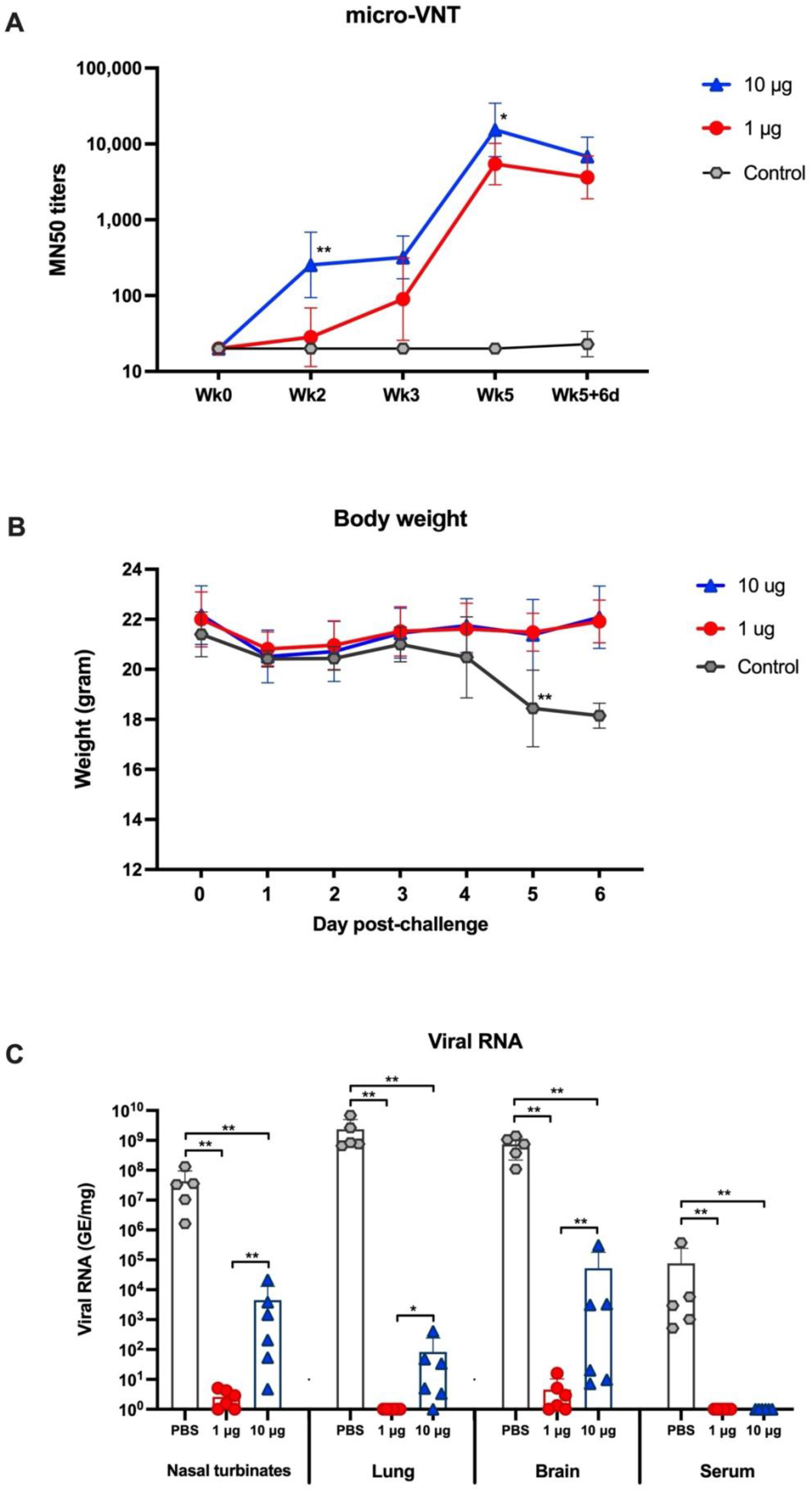
Immune response and protective efficacy results in the challenge study. (A) Kinetic responses of micro-VNT50 titer after ChulaCov19 immunization and after challenge. Data are present as GMT of micro-VNT50 titer with 95% CI. Statistical analysis was performed to compare the GMT of micro-VNT50 between 1 and 10 μg dosed mice at each time point. (B) Mean of body weight with SD after viral challenge. In the control group, 3 out of 5 mice reached euthanasia criteria on Day 5, only 2 mice were analyzed for BW on Day 6 after challenge. (C) SARS-CoV-2 viral RNA copies detected by RT-qPCR in serum and homogenized tissues of challenged animals, calculated as genomic equivalent/mg of tissue.

After SARS-CoV-2 challenge, there was no measurable decline in body weight among vaccinated groups. The average percent decline from peak to euthanasia among PBS-receiving mice was 17%. The average body weight by group from week 5 to week 5+6 days was demonstrated in the **Figure 6B**. By Day 4 after challenge, mice in the control group began to show clinical signs, first appearing in two mice with anorexia, lethargy, and rough hair coat. On Day 5, all five mice exhibited varying symptoms of increased anorexia, lethargy, immobility, rough hair coat and increased respiration rate and effort. Three out of five mice reached euthanasia criteria on Day 5, and symptoms progressed for the remaining two mice which met criteria on Day 6. In contrast, mice that received 2 doses of either 1 or 10 μg of ChulaCov19 were normal with no symptoms throughout 6 days post-challenged.

The RT-qPCR data showed that both doses of vaccine prevented the expression of SARS-CoV-2 viremia at 5 or 6 days after viral inoculation. There was no detectable viremia in mice in both high dose or low dose vaccine-treated groups while average of 7.71×10^4^ GE/mL (ranged from 1.03×10^3^ – 3.75×10^5^ GE/mL) of viral RNA was detected in PBS-received mice, **Figure 6C**. These results suggest that both dosing regimens effectively protect the mice from detectable levels of circulating virus. Moreover, the low dose regimen was also shown to induce a marked reduction in viral load in nasal turbinates, brain, and lung tissues compared to sham-treated controls. The average reduction of viral load in tissues of both vaccine-treated groups relative to the control was 99.9-100% (***Supplementary Table 1***), indicating that vaccination leads to protection against SARS-CoV-2 infection.

#### Histopathology Results in the Viral Challenge Study

In the lung, inflammation was limited to predominantly peribronchiolar proliferation of mononuclear cells, akin to an expansion of cellularity among bronchiolar lymphoid tissue but without notable follicle formation. In the nasal turbinates, all animals exhibited luminal accumulation of mucus and/or fibrin in vaccinated mice, albeit minimal to mild amounts. Immunofluorescent results mostly correlate with PCR data. No positive detection was present in the 10 μg group animals. Among the 1μg group, only one tissue had very few positive cells, the nasal epithelium. It is notable that while all mice dosed at 10-μg and 1-μg group (except 1 animal) of ChulaCov19 mRNA showed no detectable SARS-CoV-2 viral protein in tested tissues, a low level of viral mRNA was found in animals from the 10-μg group but not in the 1-μg group. In control group, positive staining is present in individual neurons of the olfactory bulb (4/4), epithelial cells of the nasal sinus (4/5), alveolar epithelial cells and macrophages in the lung (5/5), see **Table 1**.

**Table 1:** Immunofluorescent results of viral antigen in tissues

| Vaccine Dose | Mice No. | Nasal Epithelium | Olfactory Bulb | Lung | Eye, Retina |
| --- | --- | --- | --- | --- | --- |
| 1 µg | 1 | 0 | ND | 0 | ND |
|  | 2 | 0 | 0 | 0 | 0 |
|  | 3 | 0 | ND | 0 | ND |
|  | 4 | 1 | 0 | 0 | 0 |
|  | 5 | 0 | 0 | 0 | 0 |
|  | 6 | 0 | ND | 0 | ND |
| 10 µg | 1 | 0 | ND | 0 | ND |
|  | 2 | 0 | ND | 0 | ND |
|  | 3 | 0 | ND | 0 | ND |
|  | 4 | 0 | ND | 0 | ND |
|  | 5 | 0 | 0 | 0 | 0 |
|  | 6 | 0 | 0 | 0 | 0 |
| PBS | 1 | 0 | 2 | 3 | ND |
|  | 2 | 1 | 2 | 4 | 0 |
|  | 3 | 1 | 3 | 4 | 0 |
|  | 4 | 1 | 3 | 3 | 0 |
|  | 5 | 1 | ND | 4 | ND |

Note: Scores indicate subjective evaluation of the distribution of positive cells according to the following criteria: 0=negative; 1=<10% positive; 2=10-25% positive; 3=35-50% positive; 4=50-75% positive; 5=>75% positive; ND = Not Determine.

## Discussion

In this study, we described the construction and preclinical evaluation of ChulaCov19 expressing prefusion non-stabilized ectodomain of spike protein. The protein expression studies revealed S protein expressed both in intracellular and extracellular compartments when detected either by specific antibodies or patient sera (**Figure 3A**). In supernatant, we could detect both intact S and cleaved S1 and S2 at (**Figure 3B**). These results reflect the real S protein dynamic in nature as shedding of S1 could be detected in viral infection in nature (35, 36). Although several SARS-CoV-2 vaccines used an engineered S protein to abolish S1/S2 cleavage or to stabilize the prefusion stage (37–39), vaccines encoding unmodified S protein are also worth exploring as its structure is the same as native viral protein. Moreover, ChAdOx1: AZD1222 that used unmodified S has been shown to be effective (40, 41). The structural study of S protein expressed by AZ1222 showed a native-like structure mostly found in the prefusion stage. Post-translational modifications were also similar to those observed on SARS-CoV-2 (42). However, further beneficial evaluation on the use of native-like S protein structure requires in-depth analysis in clinical settings especially in immune elicitation characteristics.

Two approved mRNA vaccines, Comirnaty™ of Pfizer/BioNTech and Spikevax™ of Moderna, comprise 2 proline substitutions at residues 986 and 987 of the S-protein (known as S-2P) to stabilize the prefusion conformational structure. There has not been shown, however, that COVID-19 mRNA vaccine encoding non-stabilized spike protein is not immunogenic or is not protective against viral challenge. In these preclinical studies in mice we demonstrated that ChulaCov19, a secreted, prefusion non-stabilized ectodomain spike mRNA vaccine, elicited robust Spike-specific B-cell and T-cell responses which has also translated into efficacy in protecting transgenic mice from SARS-CoV-2 challenged.

Our immunogenicity study depicted that ChulaCov19 was highly immunogenic, in a dose-responsive relationship, even when immunized with the very low amount of 0.2 μg as measured by both live- and pseudovirus-neutralization assays. The induced NAb was highly specific to the original variant, however, cross-neutralization against the VOCs was also observed. As expected, Omicron subvariants, especially BA.4/5, showed the largest drop in micro-VNT50 titers (**Figure 4B**). This was concordant with the previous findings that Omicron subvariants could evade NAb induced by the first-generation or WT-virus-based vaccines (43). The NT50 titer decrease found in our study was similar to those of other approved vaccines as the titers against BA.1 and BA.4/5 were decreased more than 8-10 folds when compared to the WT virus (43–45). It is therefore during the surge of Omicron globally, there is a need of boosting with a first-generation vaccine or ideally with a second-generation vaccine such as a bivalent immunogen containing or encoding of Omicron’s spike protein (46, 47).

In many countries, immunization regimens have frequently employed mixtures of different vaccine platforms (also known as a heterologous prime-boost). This is especially true of the mRNA vaccines, and the approach has shown better results than homologous prime-boost with a non-mRNA-based vaccine (48). In mice, ChulaCov19 mRNA vaccine was highly immunogenic as a booster in settings primed with either inactivated or viral vector vaccine. ChulaCov19 enhanced the magnitude of both NAb and T cell responses in superior fashion to homologous 2-dose regimens of either CoronaVac or AZD1222. NT50 titers against WT and Delta variants increased 7- to 14-fold when using the heterologous approach with ChulaCov19 as compared to the homologous immunizations with CoronaVac or AZD1222 (Fig 4C). The low MN50 titer in the homologous AZD1222 group may have been influenced by anti-vector antibodies as the interval between doses was short (4 weeks). In the clinical setting, >8 weeks interval for AZD1222 was recommended to maximize the vaccine efficacy (49). However, boosting with ChulaCov19 would avoid these concerns entirely, as well as shortening the vaccination gap and increase the speed of vaccine coverage. In terms of spike-specific T cell responses, our study found that AZD1222 prime/ChulaCov19 mRNA boost induced the highest magnitude of T cell response, superior to that of all tested regimens, including the homologous mRNA vaccine (**Figure 5B**). This observation correlates with that of a recent report on a clinical study (50).

As the Omicron subvariant BA.4/5 is currently spreading worldwide, we also have assessed the cross-neutralization and found that the NAb GMT measured by psVNT50 against BA.4/5 in homologous ChulaCov19 vaccination or heterologous boosted with ChulaCov19 groups were significantly better than either of the CoronaVac or AZD1222 homologous vaccination (**Figure 4D**). This is consistent with a previous report (43).

Protective efficacy has been demonstrated in the challenge study, and there was no anamnestic antibody response detected in the ChulaCov19 mRNA vaccinated mice after viral challenge (**Figure 6A**). This is a surrogate marker indicative of vaccine effectiveness, or the sterilizing immunity as reported in the previous study (51). Moreover, the vaccinated groups showed a preventive effect on clinical symptoms, viremia, tissue viral load and the histopathology findings in hACE-2 expressing mice. Of note, no SARS-CoV-2 protein was detected in organ tissues in mice vaccinated with ChulaCov19 at either the 1 or 10 μg dose; whereas when RT-qPCR was used, low level of viral mRNA was detected in mice vaccinated with 10 μg but not 1 μg mice. Nonetheless, both dosages demonstrated a 99-100% reduction of the viral RNA in tested tissues when compared to the control group. The possible explanation of low-level detectable tissue viral RNA in the 10 μg but not 1 μg immunized mice may include the detection of free viral RNA from disintegrated virus, or a higher dose mRNA vaccine, while able to induce NAb in a dose-dependent manner, may reduce specific T-cell responses (a key immune function to eliminate cell-associated virus) as shown in the dose-response study (**Figure 5A).**

The vaccine inequity issue is a huge challenge to healthcare in LMICs. In all past pandemics, as well as the ongoing one with COVID-19, access to effective vaccines in a timely manner and has been severely limited in these countries. The most effective, COVID-19 vaccines are mRNA-based, and were first approved in the Unites Kingdom, the United States, and Europe. They were widely available in these countries for approximately a year before being accessible in other continents. LMICs received these vaccines much later and in shorter supply, as evidenced by the most recent statistic that in several African countries less than 30% of the population has received at least 1 dose of vaccine (20). Developing mRNA vaccine technology for distribution in these regions is therefore extremely important (21).

The ChulaCov19 mRNA vaccine development program has exactly this goal, striving to address the current and future pandemics in LMICs (52). The program is funded by the Government of Thailand. The promising preclinical study results presented here demonstrate that ChulaCov19 is highly immunogenic with protective efficacy, and the candidate vaccine has now been completed non-clinical toxicity and biodistribution studies and entered Phase 1 and 2 human trials. More importantly, in partnering with a domestic vaccine manufacture, BioNet Asia, the mRNA vaccine can now be manufactured and formulated locally (52). This initiative is ready to be part of the global effort to make mRNA vaccines more quickly and widely available when facing new variants or the next pandemic.

### Conclusion

This mRNA vaccine development is an effort to set up this technology platform in LMICS. Here we demonstrated that an LNP-encapsulated mRNA encoding a secreted form of prefusion non-stabilized ectodomain of SARS-CoV-2 spike protein “ChulaCov19” was able to elicit robust, specific, and protective B- and T-cell responses. With such promising results, ChulaCov19 is currently entered phase 1-2 clinical trials and manufactured locally for later clinical development. This program is a strong foundation for the next pandemic preparedness to make mRNA vaccine widely and timely accessible for LMICs including Thailand.

## Author Contribution Statement

EP, CK, DW and KR: study conception and design, EP, KT and CK: data collection, AT, AJ, KR, KP, TP, DW and KR: analysis and interpretation of results, MGA, KT, PK, NY, PP, SB, SM, TH, RIE, WW, KL and JH: reagent preparation and analysis, EP, CK and KR: draft manuscript preparation. EP, CK and KR: grant funding acquisition. All authors reviewed the results and approved the final version of the manuscript.

