## Supplementary Table 1 for "Immunogenicity and Protective Efficacy of a SARS-CoV-2 mRNA Vaccine Encoding Secreted Non-Stabilized Spike Protein in Mice"

**Supplementary Table 1:** Percent reduction of SARS-CoV-2 RNA in tissues

| **Tissues** | **% Reduction** | |
|---|---|---|
| | **1 µg/Control** | **10 µg/Control** |
| Nasal turbinates | 99.9% | 99.9% |
| Brain | 100% | 100% |
| Lung | 100% | 99.9% |

**Note:** 10 µg/Control = % Reduction of 10 µg dosed group comparing to control

1 µg/Control = % Reduction of 1 µg dosed group comparing to control
